# GORAB, a Golgi protein required for centriole structure and duplication

**DOI:** 10.1101/232272

**Authors:** Levente Kovacs, Jennifer Chao-Chu, Sandra Schneider, Marco Gottardo, George Tzolovsky, Nikola S. Dzhindzhev, Maria Giovanna Riparbelli, Giuliano Callaini, David M. Glover

## Abstract

Genome wide screens are widely believed to have identified the great majority of genes required for centriole duplication. However, seeking to clarify the partners of the *Drosophila* cartwheel protein Sas6, we identified Gorab, a known trans-Golgi associated protein that is mutated in the human wrinkly skin disease, gerodermia osteodysplastica. We now report that Gorab is present not only in the trans-Golgi but also in association with Sas6 at the core of the centriole. Flies lacking Gorab show defects in centriole duplication in many tissues and are also uncoordinated due to basal body defects in sensory cilia, which lose their 9-fold symmetry. We demonstrate the separation of centriole and Golgi functions of *Drosophila* Gorab in two ways: First, we have created Gorab variants that are unable to localize to trans-Golgi but can still rescue the centriole and cilia defects of *gorab* null flies. Secondly, we show that expression of C-terminally tagged Gorab disrupts Golgi functions in cytokinesis of male meiosis, a dominant phenotype overcome by a second mutation preventing Golgi targeting. We discuss the tissue specific requirement of Gorab for centriole duplication in the context of its split functions.

## Introduction

Centrioles are found at the core of centrosomes, where they are required for the fidelity of cell division, and at the base of cilia, required for processes as diverse as motility and cell signalling. Thus, their malfunction is associated with a wide range of diseases from cancer to ciliopathies and microcephaly. Key components of the conserved canonical pathway for centriole duplication were first identified through genetic studies in *C.elegans*^1^ and genome-wide RNAi screens in *Drosophila*^2,3^. Centriole duplication requires Polo-like kinase 4, which phosphorylates the centriole cartwheel protein, Ana2/STIL, to enable it to physically interact with Sas6 at the site of procentriole formation ^4,5^ The resulting 9-fold symmetrical structure reflects an inherent symmetry of 9 interacting dimers of Sas6in the cartwheel at the proximal part of the centriole. Cep135 (Bld10) is also required for cartwheel formation in *Chlamydomonas* and *Paramecium*^6,7^and human Cep135 links Sas6 to Sas4 (CPAP)^8^, which is associated with and is required to promote the polymerisation of centriolar microtubules (MTs)^9–11^. Although the newly assembled centriole reaches its full length in early mitosis, it cannot duplicate or organize pericentriolar material (PCM) until it has passed through mitosis^12,13^. This requires Cep135 to establish another molecular network by recruiting Anal in *Drosophila* or its human counterpart, Cep295,and in turn Asterless/Cep152and pericentrin-like protein^14^. This process, known as centriole to centrosome conversion enables the daughter centriole to recruit Plk4, as a partner of Asterless/Cep152, as well as components of the peri-centriolar material which organises cytoplasmic MTs.

In the interphase cell, the centrosome is localized in the vicinity of the Golgi apparatus, which lies at the heart of the secretory pathway and is a key interchange for vesicular trafficking. The positioning of the Golgi and the trafficking of vesicles rely on cytoplasmic MTs, which have been shown to organize by the Golgi through AKAP450 (counterpart of pericentrin), the *cis*-Golgi protein GM130, the *trans*-Golgi CLASPs (cytoplasmic linker-associated proteins), and additional proteins in a multiprotein complex^15–17^. Interphase cells can also utilise the mother centriole to template the formation of cilia. The Golgi is also known to participate in this process by providing membrane for the cilium and the ciliary pocket close to the mother centriole^18–20^. In mouse and human cells, this requires the golgin GMAP-210, which interacts with IFT-20, part of the intraflagellar transport complex required for ciliogenesis ^21,22^. These and other interactions between the centrosome and Golgi are only just beginning to be explored and require further study.

Although 9 dimers of Sas6 adopt the 9-fold symmetrical structure of the cartwheel, Sas6 alone cannot confer 9-fold symmetry on the centriole; mutations that destroy the symmetry of Sas6’s interactions *in vitro*^23^or that prevent Sas6 self-oligomerisation are still able to make 9-fold symmetrical centrioles ^24^ This suggests that additional, yet unknown factors must contribute to the establishment of 9-fold symmetry, raising the question of whether Sas6-interacting proteins had been missed by earlier genome-wide screens. This led us to use proteomic approaches to search for additional componentsof the *Drosophila* centriole, of which we now describe a new Sas6 partner, the fly counterpart of the Golgi-associated protein Gorab. Human GORAB is mutated in gerodermia osteodysplastica, a disease characterised by wrinkly, non-elastic skin and osteoporosis^25^. GORAB also interacts with the kinase-like protein SCYL1 ^26^ that participates in the Golgi-ER trafficking of COPI vesicles ^27,28^. GORAB localises to the trans-Golgi compartment from which it is rapidly displaced by Brefeldin A suggesting its membrane association ^29^. We now show that null mutants of *Drosophila Gorab* have greatly reduced numbers of centrosomes in embryos and diploid larval tissues and have ciliary defects in neurosensory organs that include defects in 9-fold symmetry, and which result in loss of coordination. We created a missense and a small deletion mutant of Drosophila Gorab, both of which are no longer able to localise to the Golgi but are still able to rescue centriolar phenotypes of a *gorab* null. We also generate a dominant negative form of GORAB whose cytokinesis phenotypes depends upon its localisation to the Golgi. Together these findings indicate Gorab to have dual roles, at the Golgi and at the centriole, suggesting the possibility that some of the broad range of phenotypes observed in gerodermia osteodysplastica patients might be associated with additional defects in centrioles and/or cilia.

## RESULTS

### Gorab copurifies with Sas6 and is found at centrosomes and Golgi

To gain further insight into the mechanism of centriole duplication, we wished to identify partners of Sas6, one of the first proteins detected at the site of procentriole formation. To this end, we affinity purified Sas6 and associated proteins from syncytial embryos of transgenic *Drosophila* expressing GFP-tagged Sas6 and from cultured cells after inducible expression of Protein-A-tagged Sas6. Surprisingly, we found no significant enrichment of other centriole proteins in Sas6 pull-downs from embryos but we consistently identified large numbers of peptides derived from the *CG33052* gene product (Fig. 1a; Table S1). This complex persisted following treatment with high salt (440mM NaCl) and was not affected by inhibition of protein phosphatases with okadaic acid suggesting it is insensitive to the protein phosphorylation state. Similarly, mass spectrometry of Sas6 complexes purified from cultured cells consistently identified CG33052 of significance excluding proteins commonly found in control purifications with affinity beads alone. BLAST searches identified CG33052 as the counterpart of human GORAB, which is mutated in the inherited disease Gerodermia Osteodysplastica, leading us to name the *Drosophila* CG33052 gene *gorab*.

**Fig. 1.**
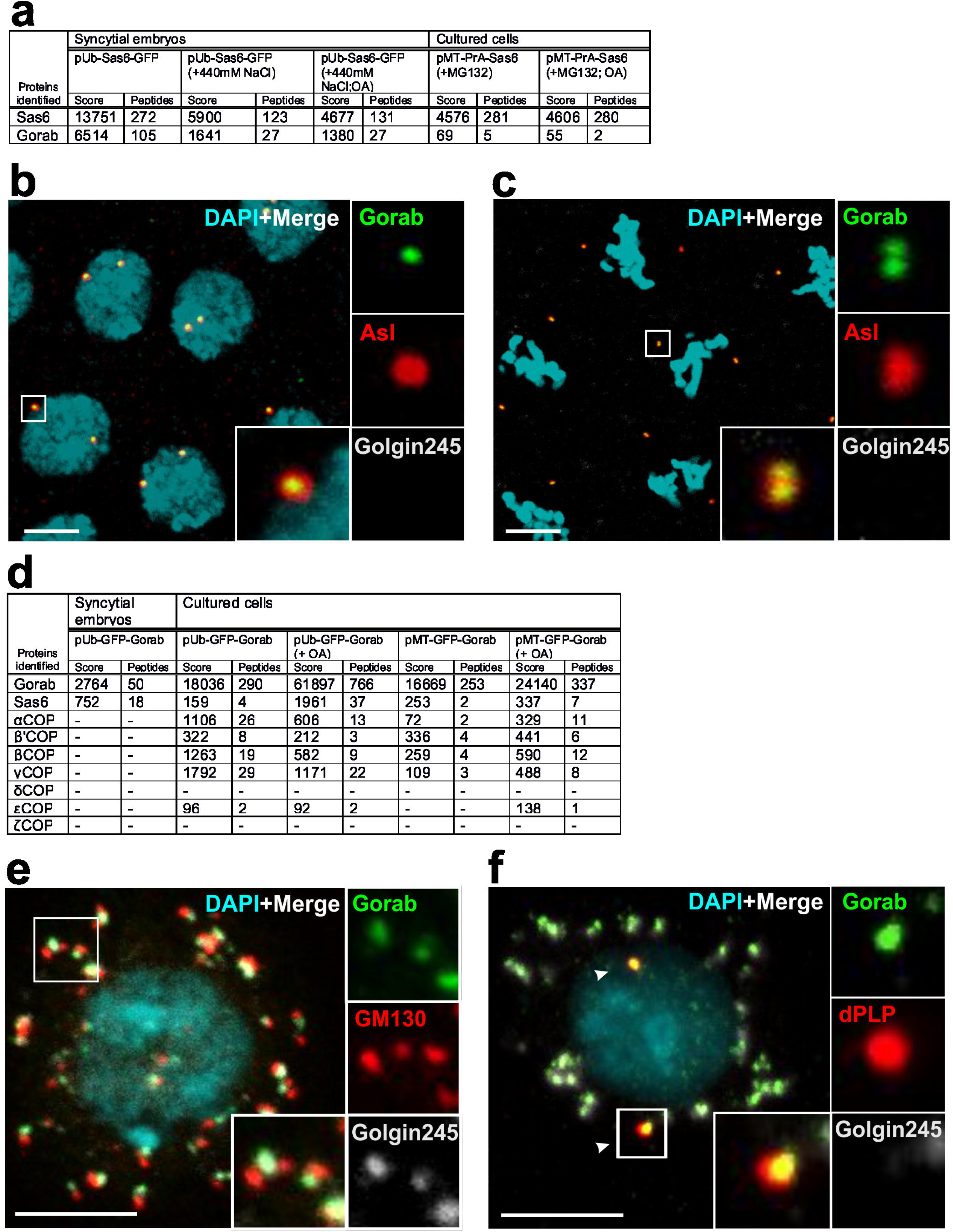
Gorab associates with both Centrosomes and Golgi. (**a**) Co-purification of Gorab with GFP-Sas6 from polyUbiquitin-Sas6-GFP *Drosophila* embryos or Protein A-Sas6 expressed from metallothionein promoter in D-Mel cells. Extracts made in isotonic or high salt (440mM NaCl) buffer and with okadaic acid (OA) and MG132 as indicated (Materials and Methods). Proteins commonly identified in control purifications of GFP or other GFP-tagged proteins are excluded but given in tables S1A-E. Scores (Mascot) and numbers of peptides detected by mass spectrometry are indicated. (**b, c**) Syncytial embryos expressing poly-Ubiquitin-GFP-Gorab and stained to reveal Asterless (Asl) and the trans-Golgi Golgin245 in a field of interphase (**b**) and mitotic (**b**) nuclei. Insets show 5x enlargment of indicated region of interest (ROI). Scale bar: 5 μm. (**d**) Affinity purification of tagged Gorab from poly-Ubiquitin-GFP-Gorab Drosophila embryos or D-Mel cells stably transformed with poly-Ubiquitin-GFP-Gorab or p-metallothionein (pMT)-GFP-Gorab (induced with 1mM CuSO4 for 22h). Co-purifying Sas6 and COPI complex proteins selected from the full list of co-purified proteins in tables S2 A-E. (**e**) Cultured D-Mel cells immuno-stained to reveal Gorab, GM130 (cis-Golgi), and Golgin245 (trans-Golgi). Insets show 3x enlargement of ROI. Scale bar: 5 μm. (**f**) Cultured D-Mel cells immunostained to reveal Gorab, dPLP (centrosome), and Golgin245 (trans-Golgi). Arrowheads indicate centrosomes. Insets show 3x enlargement of ROI. Scale bar: 5 μm.

To further confirm that Sas6 and Gorab copurify as a complex, we established transgenic fly and cell lines to express GFP-tagged Gorab. When we purified GFP-Gorab from syncytial embryos of transgenic flies, we consistently co-purified Sas6 as its only centriole associated partner (Fig.1d; Table S2). Sas6 was also a consistent Gorab partner in purifications from cultured cells, irrespective of whether GFP-Gorab was expressed constitutively or from the inducible metallothionein promoter. However, in addition to Sas6, the list of interactors also contained the αCOP, β’COP and εCOP subunits of the cage-like subcomplex of the COPI coatomer complex and the γCOP and βCOP subunits of its adaptor subcomplex (Fig. 1d; Table S2). We did not detect any COPI complex proteins in control purifications of GFP expressed in cultured cells or in purifications of Sas6. Nor did we detect COPI proteins associated with Gorab purified from syncytial embryos (Fig. 1d; Table S2), a stage prior to the onset of Golgi assembly ^30–32^.

The repeated copurification of Gorab with Sas6 suggested an association with the centrosome. Indeed, when we examined syncytial blastoderm embryos expressing Ubq-GFP-Gorab transgenes, we found that GFP-Gorab localised exclusively to centrosomes revealed by anti-Asterless counter staining in interphase (Fig. 1b) and mitosis (Fig. 1c). As the Golgi apparatus is not properly assembled at this stage, we could not detect staining of the trans-Golgi Golgin245. Thus, at least in the absence of the Golgi, Gorab could be a *bone* fide centrosome component accounting for its physical association with Sas6.

We then sought to determine the sub-cellular localisation of Gorab in cell types with established Golgi apparati. We therefore raised antibodies against Gorab (Material and Methods) and used them to reveal Gorab in cultured D-Mel2 cells relative to markers of the cis-(GM130) and trans-Golgi (Golgin 245). We found the greater part of all Gorab in the cell was associated with the trans-Golgi apparatus, as also found for its human counterpart ^29^ (Fig. 1e). However, co-staining showed that Gorab was also present in dPLP associated punctae indicating its presence at the centrosome independent of Golgin 245 (Fig. 1f). Moreover, in mitosis, when Golgi components become dispersed throughout the cell, we found Gorab persisted at the centrosome (Fig. S1a). We made similar findings when examining the localisation of Gorab in cells of larval wing imaginal discs and in the larval central nervous system of animals ubiquitously expressing GFP-Gorab (Fig. S1b and c). Gorab was present in an abundant and rather dynamic pool situated on trans-Golgi that becomes dispersed during mitosis and a smaller fraction that is stably associated with the centrosome in both interphase and mitotic cells in these tissues.

### Gorab directly binds Sas6 at the centriole core

The co-purification of Gorab and Sas6 and their colocalisation in centrosomes raised the question of whether they were direct binding partners. To address this, we expressed and purified GST-tagged Gorab from E.coli on Glutathione Sepharose 4B resin. When such beads were incubated with ^35^S-Met-labelled full-length Sas6, synthesised by coupled transcription-translation, we could detect binding of Sas6 to immobilized Gorab (Fig. 2a). To narrow down the region of interaction, we made a series of N- and C-terminally truncated S^35^-labelled Sas6 peptides for similar binding experiments (Fig. 2b; Fig. S2). Together these experiments identified a region of Sas6 between 351 and 462 amino-acids that could interact directly with Gorab.

**Fig. 2.**
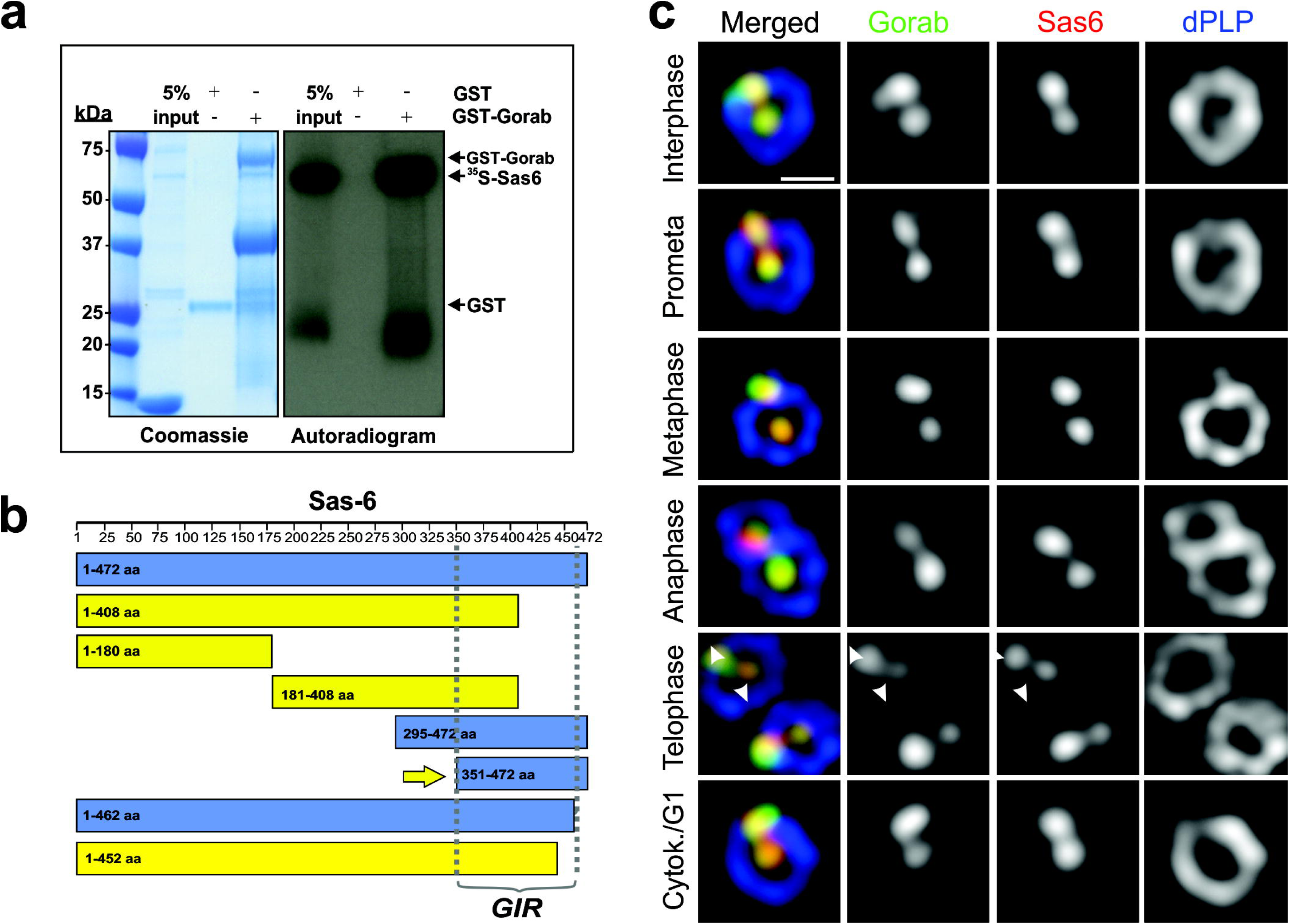
Gorab directly interacts and colocalises with Sas6. (**a**)*In vitro* assay of the Gorab-Sas6 interaction. GST-Gorab was incubated with ^35^S-Methionine-labelled Sas6 and resulting complexes subjected to SDS-PAGE and autoradiography. (**b**) Schematic of Sas6 protein fragments tested for Gorab interaction. Blue, Sas6 fragments interacting with Gorab. Yellow, Sas6 fragments not interacting with Gorab. Arrow indicates minimal interacting fragment (GIR: Gorab Interacting Region). (**c**) 3D-SIM localization of endogenous Gorab (green) and Sas6 (red) throughout the cell cycle in D.Mel-2 cells relative to zone III marker, dPLP (blue). Arrowheads indicate site of procentriole formation. Scale bar, 250 nm.

The direct binding of Gorab to Sas6 together with its centrosomal localization suggested it should be associated with the centriole. To address this, we used structured illumination microscopy (SIM) to reveal Gorab in relation to Sas6, which occupies zone I at the core of the centriole, and dPLP, which forms a ring surrounding the mother centriole throughout the whole cell cycle. This dPLP ring is only completed around the daughter centriole during passage through mitosis in centriole to centrosome conversion. We found that Gorab and Sas6 are both present in zone I of the mother and daughter centriole throughout the centrosome cycle. Moreover, once centriole to centrosome conversion has been completed allowing mother and daughter to disengage in telophase, Sas6 and Gorab are both recruited to the site of procentriole formation (arrowheads in Fig. 2c). Thus Gorab and Sas6 associate at the core of the centriole from the very onset of its duplication.

### *Gorab^null^*-derived embryos display mitotic defects due to centrosome loss

The above findings suggested a potentially new dimension to the roles of this Golgi-associated protein that we addressed by generating null mutants of *Drosophila gorab* to study the effects of loss of the protein upon development. We used CRISPR/Cas9 mutagenesis to simultaneously target the 5′ and 3′ ends and exon1 of the *gorab* gene (Methods) and generated 475 and 1168 bp deletion mutants (*gorab^1^* and *gorab^2^*), each of which eliminated a significant part of the coding sequence including the ATG initiation codon (Fig.3a). We were unable to detect any Gorab protein in Western blots of whole fly extracts (Fig. 3c) in homozygotes of either allele leading us to consider both mutant alleles to be nulls.

**Fig. 3.**
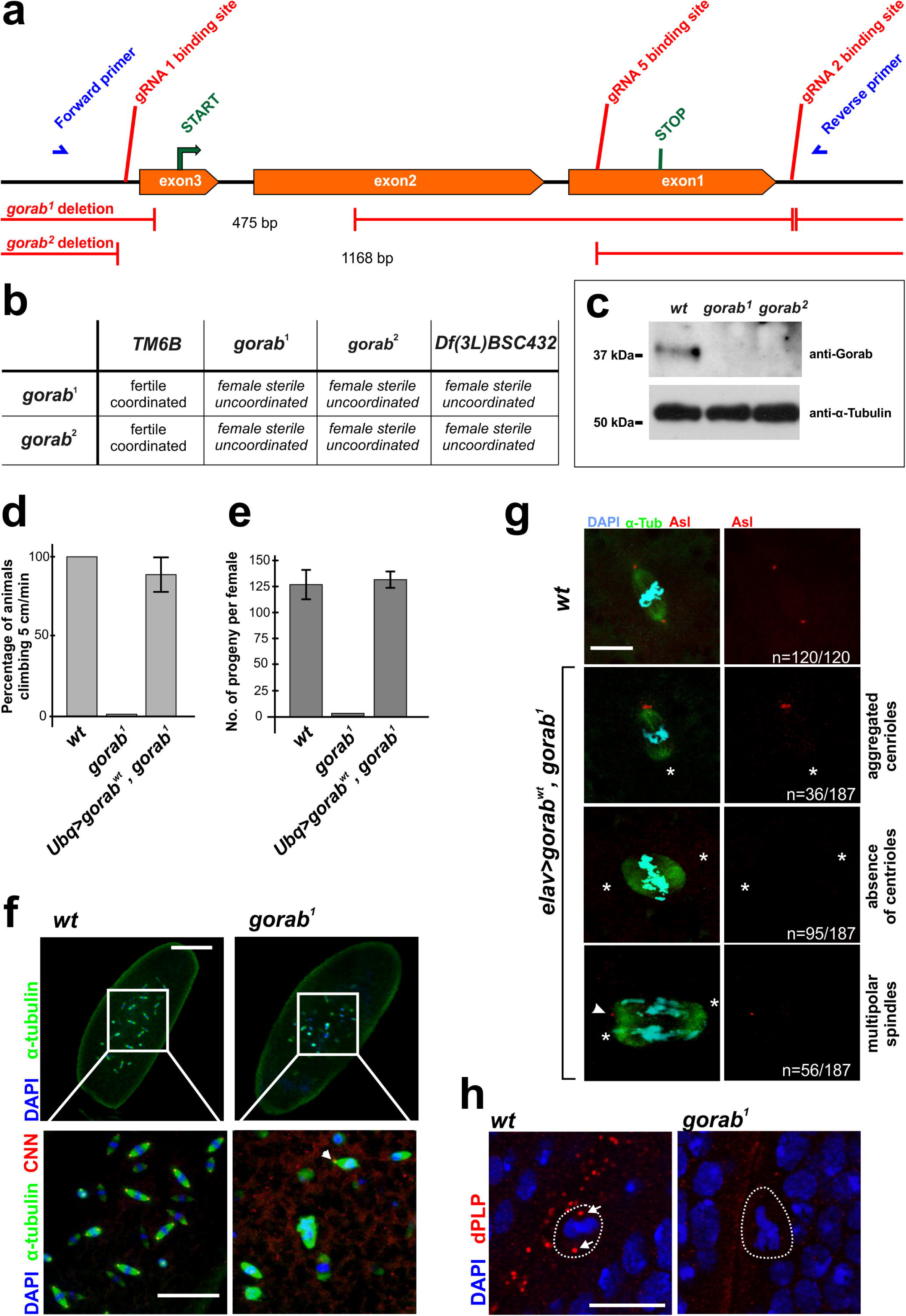
*gorab* null mutant flies show female sterility and coordination defects. (**a**) Schematic of *gorab* gene showing guide RNA binding sites (red) for CRISPR/Cas9 mutagenesis and primer binding sites for detection and sequencing of indicated *gorab^1^* and *gorab^2^* deletion mutants (blue). (**b**) Complementation tests for female fertility and coordination. 200 individuals tested per genotype. (**c**) Western blot of extracts from 5-5 newly eclosed females of indicated genotypes. Gorab revealed by antibody raised against its N terminal fragment. α-tubulin, loading control. (**d**) Climbing ability of wild type, *gorab^1^*and rescued (N-terminal-GFP-tagged *gorab^wt^* cDNA expressed from ubiqutin promoter in *gorab^1^*background); flies raised at 29 °C. Cohorts of 15 flies and scored for number climbing 5 cmin 1 min. Means with s.d.(bars); 3 independent experiments (n=45). (**e**) Fertility of wild type, *gorab^1^*and rescued (as in **d**) females individually mated with wild type males at 25°C over 6 days. Means with s.d. (bars); 15 females per genotype. (**f**) Embryos from wild-type and *gorab^1^*mutant mothers stained to reveal α-tubulin (green), centrosomin (CNN, red), and DNA (blue). Lower panels show 3X magnification of highlighted area; Arrowhead, single centrosome of monopolar spindle. Scale bar, 100 μm (upper panel); 35 μm (lower panel). (**g**) Embryos from *elav>gorab^WT^,gorab^1^* mothers stained to reveal α-tubulin (green), asterless (Asl, red), and DNA (blue). Arrowhead, third pole of a multipolar spindle; asterisks, spindle poles lacking centrioles. Scale bar, 10 μm. (**h**) Wing discs from wild type and *gorab^1^* larvae immunostained against dPLP to reveal centrosomes (red).Dashed lines highlight a mitotic cell. Arrows point to centrosomes of the mitotic cells. Scale bar: 10 μm.

Surprisingly, both homozygous *gorab^1^* and *gorab^2^* animals developed to adulthood. However, the flies that emerged at 25°C moved slowly and were uncoordinated; when raised at 29°C, they were entirely unable to climb or fly (Fig. 3d, video S1). Furthermore, embryos derived from *gorab* mutant females raised at 25 °C failed to develop. This maternal effect lethality was evident even when homozygous mutant mothers were mated to wild-type males (Fig. 3e). In contrast, there was no reduction in the fertility of *gorab^1^* males suggesting the mutation did not affect spermatogenesis. A similar extent of female sterility and loss of coordination was also observed in flies trans-heterozygous for the two alleles, or when either mutant allele was heterozygous to a large chromosomal deficiency, suggesting that these defects result exclusively from the loss of Gorab function (Fig.3b). Accordingly, female sterility and loss of coordination were completely rescued when we ubiquitously expressed a wild-type *gorab* transgene in the *gorab^1^*mutant background (Fig. 3d, e). Thus, the primary consequences of the loss of Gorab function are female sterility and loss of coordination.

Coordination defects have previously been attributed to defective neurosensory cilia of the chordotonal organs ^33,34^ Accordingly, we could rescue the coordination defects but not the female sterility of *gorab^1^*flies by expressing *UAS-gorab* only in the nervous system using the pan-neural driver *elav-GAL4.* This allowed us to recover properly coordinated adult females from which we collected sufficient embryos to analyse their development. Wild-type syncytial embryos undertake 13 rapid rounds of synchronous nuclear division cycles in a shared cytoplasm over a period of 2 hours. In contrast, only 50% of *gorab^1^*-derived embryos showed any nuclear division cycles and these did not proceed beyond 5 or 6 rounds. Centrosomes, revealed by anti-centrosomin (Cnn) or anti-Asterless (Asl) staining were dramatically reduced in number in *gorab^1^*-derived embryos and were completely absent from the poles of the majority of mitotic spindles (Fig. 3f, g). The extensive disorganisation of *gorab^1^*-derived embryos is in line with the known requirement for centrosomes to progress through the syncytial cycles. These observations highlight a requirement for maternally provided Gorab for centrosome duplication and thereby the nuclear division cycles of the embryo. We also observed the loss of centrosomes in the imaginal discs of *gorab^1^* homozygous larvae indicating a requirement for Gorab for centrosome duplication in other diploid tissues (Fig. 3h).

### Coordination defects of *gorab* flies reflect absence of daughter centrioles and loss of 9-fold symmetry in ciliary organs

The severe loss of coordination that arise in flies defective for centriole duplication reflects abnormal function of femoral chordotonal organs (fChOs) of the adult legs whose ciliated neurons have basal bodies derived from centrioles (Fig. 4a). In the wild-type fChO, immunostaining reveals *Drosophila* pericentrin-like protein (dPLP) in mother and daughter centriole-derived structures, of which the mother forms the basal body of the cilium. Transgenic GFP-rootletin, ciliary coiled-coil protein, identifies the ciliary rootlet connecting the basal bodies to the cell body and phalloidin staining reveals actin in the scolopale cell that envelopes two cilia. We found that the fChOs of *gorab^1^* mutant flies had highly disorganised ciliary rootlets, many of which were disconnected from basal bodies (Fig. 4a,b). The dPLP associated structures were also disorganized in *gorab^1^* mutants but the presence of scolopale rods indicated that ciliary structures are still present (Fig. 4a, b).

**Fig.4.**
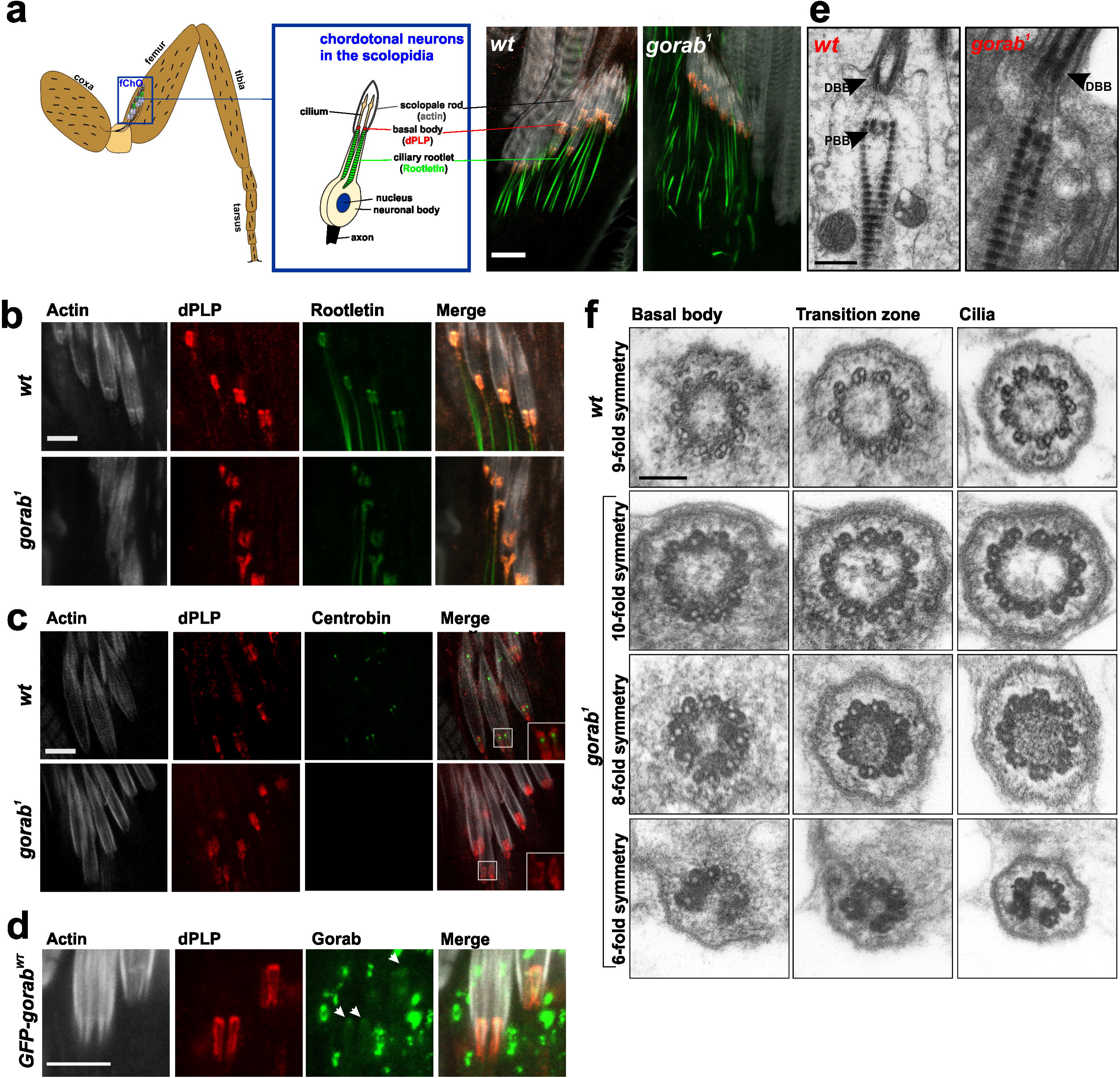
Loss of daughter centriole and asymmetrical mother centrioles in *gorab* mutant ciliated neurons. (**a**)Schematic of fermoral chordotonal organs (fChO) and their imaging in wild type and *gorab^1^* mutant to reveal transgenic Rootletin-GFP (green), dPLP in basal bodies (red), and actin in scolopale rods (white). Scale bar,10 μm. (**b**) Detail of basal body organisation in wild-type and *gorab^1^* stained as in A. Bar,10 μm. (**c**) Localisation of daughter centriole specific YFP-Centrobin (Cnb, green) in wild-type and *gorab^1^fChO* basal bodies also stained to reveal dPLP (red) and Actin (grey). Scale bar, 10 μm. (**d**) Localization of GFP-tagged Gorab expressed in *gorab^1^* mutant background. Arrowheads, GFP-Gorab at basal bodies. Scale bar, 10 μm. (**e**) EM images of longitudinal sections wild-type and *gorab^1^*fChOs. Arrowheads, distal (DBB) and proximal (PBB) basal bodies. Scale bar, 0.2 μm. (**f**) Transverse sections of basal bodies, transition zones and cilia in wild-type and *gorab^1^*imaged by TEM. Scale bar, 0.1 μm.

The two basal bodies of the fChOs can be distinguished because only the proximal one (corresponding to a daughter centriole) has associated Centrobin ^35^. However, we found that 80% of basal bodies in ciliated cells of *gorab^1^* flies had no Centrobin (Fig. 4c). Only 20% showed some anti-Centrobin positive dots that stained more weakly than in wild type. This suggests a failure of centriole duplication prior to basal body formation. We confirmed that GFP-Gorab could restore the correct number of basal bodies and the organisation of the fChOs of *gorab^1^* mutants (Fig. 4d).

To determine whether there were other abnormalities of the basal bodies of *gorab^1^* ciliated neurons we examined them in longitudinal and transverse section by electron microscopy (EM) (Fig. 4e, f). Longitudinal sections of *gorab^1^*chordotonal organs revealed an absence or reduction of proximal basal bodies in accord with the lack of Centrobin signal (Fig. 4e). However, mother centrioles were always present all distal basal bodies investigated (n=11). Transverse section of the *gorab^1^* ciliated cells revealed striking defects in radial symmetry of the remaining mother centrioles of the basal body (Fig. 4f). Only 3 of the 16 centrioles observed showed a normal 9-fold symmetrical arrangement of the microtubules, whilst the rest had symmetries varying between 6-fold (n=5/16), 8-fold (n=6/16) and 10-fold (n=2/16). The defective symmetry extended from the basal body through the transition zone to the distal part of the cilium (Fig.4f). This phenotype is strikingly similar to the defects in centriole symmetry seen in Drosophila Sas6 mutant animals^36^.

Thus, the loss of coordination in *gorab* mutants reflects the abnormal anatomy of the fChO resulting from the loss of daughter centrioles and the abnormal symmetry of the remaining mother that constitutes the basal body.

### Golgi and Centriole localisation domains of Gorab overlap

To address Gorab’s functions at the centrosome and at the Golgi, we first wished to define functional domains within the protein responsible for its localisation to these two sites. We began by defining the domain within Gorab responsible for binding Sas6. To this end we immobilised GST-tagged Sas6 on Glutathione Sepharose 4B beads and assessed its ability to bind^35^ S-Met-labeled Gorab and or its fragments synthesised *in vitro* (Fig. 5a,b; Fig. S3a). We narrowed down a strong direct interaction of Sas6 to an interval of Gorab between amino-acids 191 and 318. A Gorab fragment of amino-acids 244 and 338 also bound but more weakly (Fig. S3a). We then found that three non-overlapping smaller deletions within this region of Gorab (aa260-266, aa267-281 and aa282-286) still permitted Sas6 binding whereas a 27aa deletion (aa260-286) abolished the interaction, thus defining a Sas6-interacting-domain (SID) within this last interval (Fig. 5c,d) To confirm that the *in vitro* interaction could reflect a similar interaction *in vivo,* we transiently transfected cultured *Drosophila* cells with myc-tagged Sas6 and either full length Gorab or Gorab^ΔSID^ tagged with GFP. Western blots of GFP-pulldowns from extracts of these cells showed that Sas6 interacted with Gorab but not Gorab^ΔSID^ (Fig. S3b) confirming the *in vitro* binding results.

**Fig. 5.**
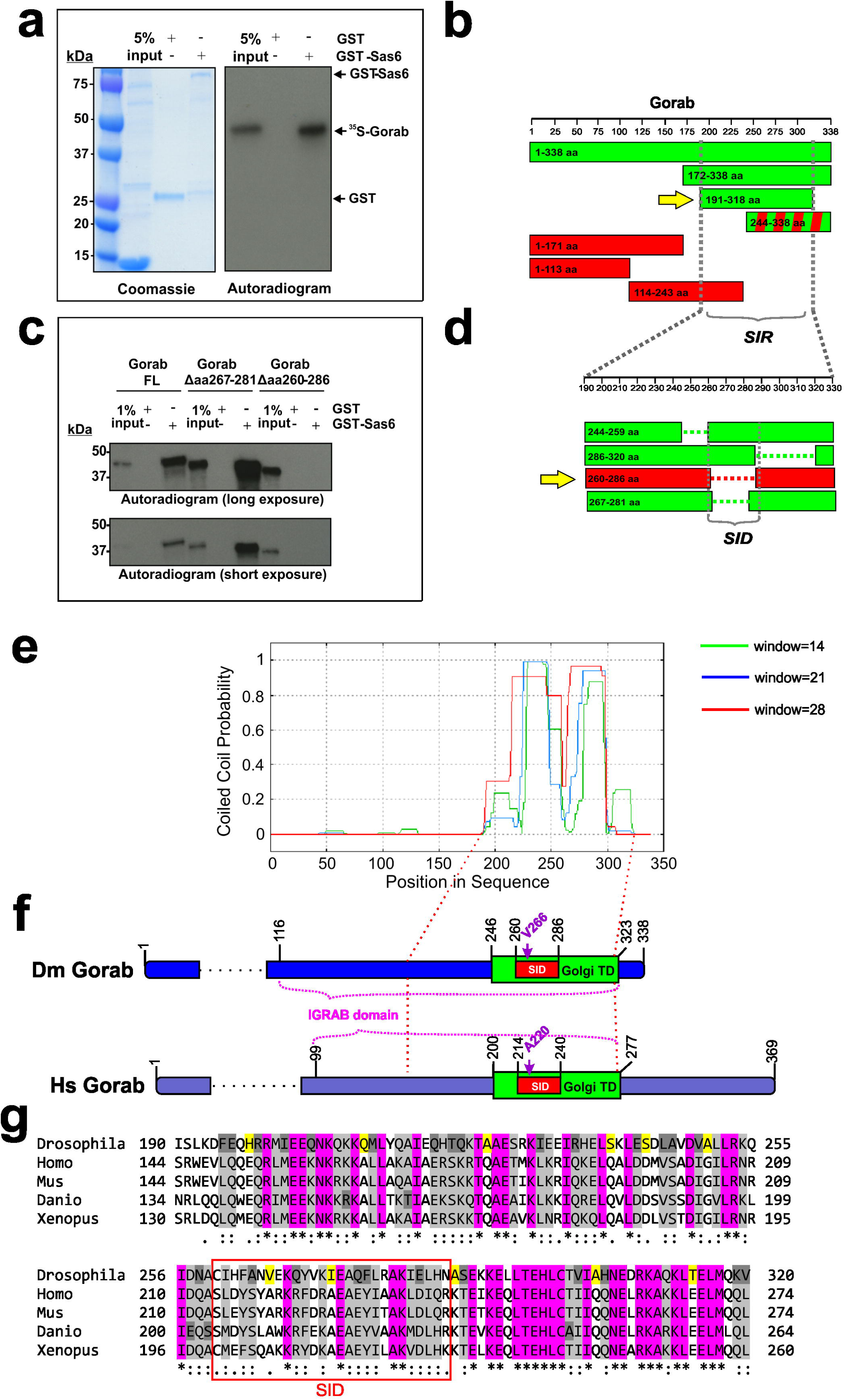
Domain structure of Gorab. (**a**) *In vitro* interaction of GST-Sas6 and ^35^S-Methionine-labelled Gorab. (**b**) Identification of minimal Sas6 interacting region (SIR) in Gorab. Green bars, Gorab fragments interacting with Sas6. Red bars, Gorab fragments not interacting with Sas6. Green and red stripes, weak interaction. Arrow indicates minimal interacting fragment. (**c**) Sas6 interaction with ^35^S-Methionine-labelled full length (FL) Gorab and indicated deletion variants. (**d**)Minimal Gorab deletion that abolishes Sas6 interaction. Green bars, Sas6 interacting Gorab constructs; red bars, non-interacting constructs. Arrow indicates minimal deletion abolishing the interaction with Sas6, the Sas6 interacting domain (SID). (**e**) Prediction of coiled-coil region in Gorab, by Coils server by scanning windows of 14, 21 and 28 residues. (**f**) Comparison of domain topologies of *Drosophila* and human Gorab. Sas6 interacting domain (SID, red), Golgi targeting domain (Goldi TD, green), and IGRAB domain are indicated. (**g**) Alignment of predicted coiled-coil region of *Drosophila* Gorab and five vertebrate species homologues. Pink, amino acids conserved between all; grey, similar amino acid groups; dark grey, single divergent amino acids; and yellow, a single divergent amino acid in *Drosophila.* Alignment generated with Clustal Omega. Red box, SID.

The SID domain lies within a predicted coiled-coil region (approximately amino-acids 190 to 320; Fig. 5e) that lies in an analogous position to the predicted coiled-coil domains of human GORAB (Fig. 5f). Previous studies of human GORAB defined a minimal tested fragment able to bind Arf5 and Rab6 in a yeast 2-hybrid system and in pull-down experiments; the IGRAB domain, amino-acids 99-277 ^29^. These same workers defined a minimum fragment tested for abily to localise to the Golgi; the Golgi Targeting Domain (GTD), amino-acids 200-277 ^29^ that corresponds to amino-acids 246-323in Drosophila Gorab. Previous studies of gerodermia osteodysplastica patients have identified a missense mutation within the GTD, Ala220Pro, that is sufficient to disrupt Golgi localisation ^29,37^ Sequence comparison of the *Drosophila* and human GORAB proteins indicated they were 70% similar and 40% identical over the GTD (Fig. 5g). Within the GTD of *Drosophila* Gorab we could identify a conservative amino acid change to Valine at position 266 corresponding to Ala220 in the human GORAB sequence, the site of the Golgi mislocalising mutation. Thus, curiously, the SID, required for Sas6 binding, lies within the GTD and the single amino-acid required for Golgi targeting mutation lies within the SID. Thus the parts of Gorab required for Golgi and centrosome localisation overlap.

### Gorab variants unable to localise to Golgi retain centriolar function

The overlapping nature of the potential site for Golgi localisation and Sas6 interaction led us to ask whether mutations in this region would necessarily affect both properties. Could we, by modelling the missense mutation affecting GORAB’s Golgi localisation in gerodermia osteodysplastica patients, disrupt the Golgi localization of Drosophila Gorab, and if so, how this might affect the centrosomal localisation and function of the protein. We therefore generated a *Drosophila gorab* transgene with a Val266Pro mutation to mimic the human Ala220Pro mutant together with transgenes having deletions corresponding to the SID, the GTD, an N-terminal part of the GTD (N-GTD), and a C-terminal part (C-GTD) (Fig. 6a). We then generated transgenic flies with these *gorab* variants under the control of constitutive or inducible promoters integrated into the same genomic location using a site-specific integrase system so that they would each be expressed at a similar level (Materials and Methods). An eGFP tag on the N-termini of the mutant transgenes allowed us to determine the subcellular localisation of the gene products in cells of larval imaginal discs. In agreement with our earlier findings, the vast majority of wildtype eGFP-Gorab colocalized with Golgin245 trans-Golgi marker and a small fraction was recruited to centrosomes. By contrast, the mutant forms of Gorab lacking the SID, the GTD and the N-terminal part of the GTD were diffusely distributed throughout the cytoplasm and had no specific accumulation at either Golgi or centrosomes (Fig. 6b). However, both the V266P missense mutant and C-terminal GTD deletion retained their centrosomal localization despite losing their ability to localise to the Golgi (Fig. 6b).

**Fig. 6.**
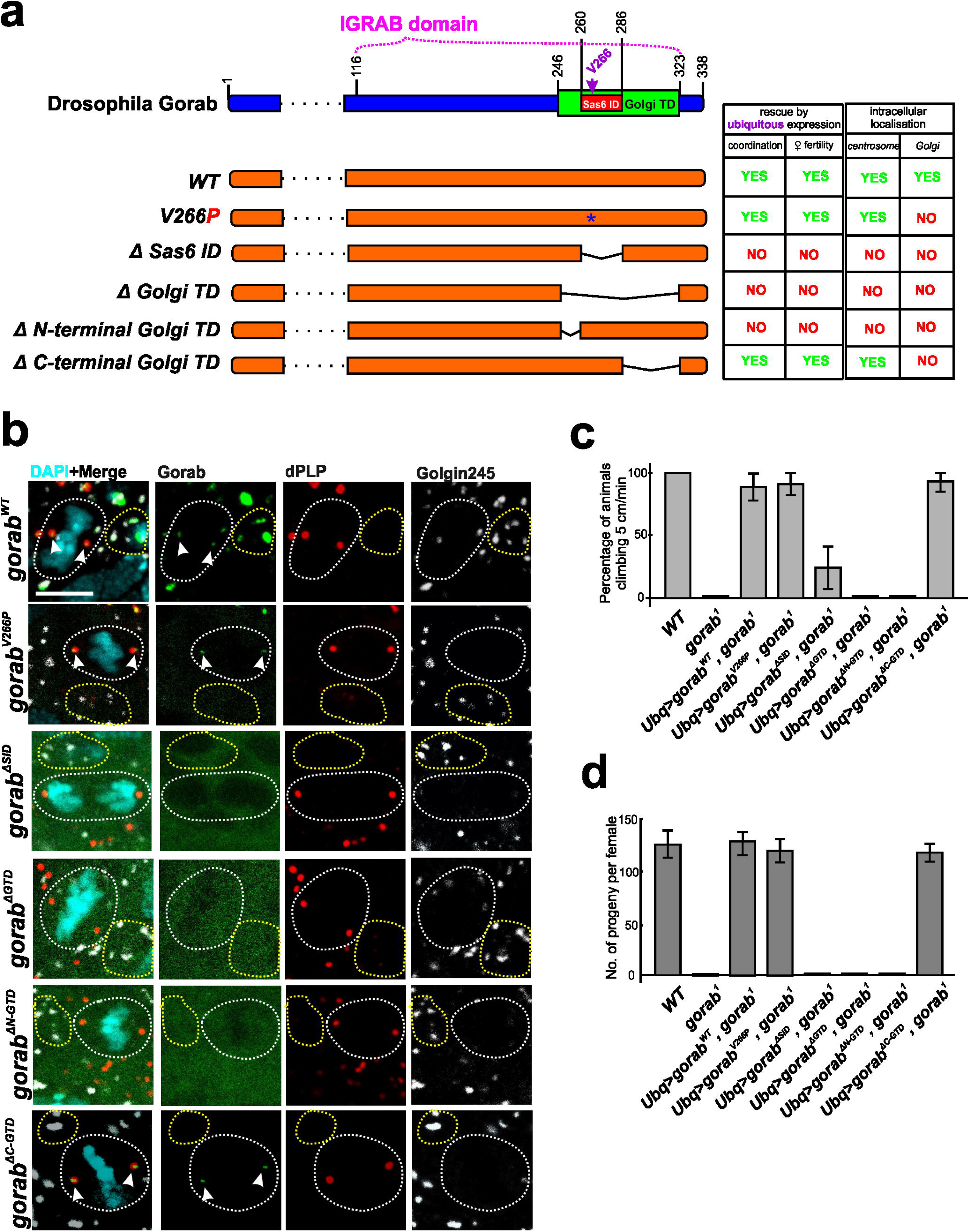
Functional Domains of Gorab. (a) Schematic of mutant Gorab transgenes used in localization and rescue experiments. Asterisk, position of Val266 to Pro substitution. Gaps connected with thin line indicate extents of deletions. Tables summarizes rescue and localization experiments alongside indicated mutant forms. (b) Intracellular localization of Gorab mutant transgenes, Wing discs from larvae expressing the indicated GFP-tagged Gorab mutant protein stained to reveal dPLP (red) and Golgin245 (grey). White dashed lines highlight mitotic cells, yellow dashed lines indicates interphase cells with assembled Golgi. Arrowheads, centrosomes of mitotic cells. Scale bar, 5μm. (**c**) Rescue of climbing ability by ubiquitous expression of indicated N-terminally GFP-tagged transgene in *gorab^1^* flies raised at 29 °C. Cohorts of 15 flies observed for 1 minute; flies climbing 5 cm recorded. Means with s.d. for 3 independent experiments (n=45). (**d**) Rescue of female sterility by ubiquitous expression of indicated N-terminally GFP-tagged transgenes in *gorab^1^* background. Individual females mated with wild type males and left at 25 °C for 6 days. Number of progeny quantified. Means with s.d. for 15 females/ genotype.

The centrosomal localization of V266P and ΔC-GTD mutant Gorab raised the question whether these variants are still functional at the centrosomes. To address this, we introduced these *gorab* mutant transgenes into a *gorab^1^* null mutant background and found that the *gorab^V266P^* and *gorab^ΔC-GTD^* transgenes restored the climbing ability of *gorab^1^*flies raised at 29 °C to a similar extent as *gorab^WT^*whereas *gorab^ΔGTD^*, *gorab^ΔN-GTD^*and *gorab^ΔSID^* failed to do so (Fig. 6c).In addition, *gorab^V266P^* was also shown to localize to the basal bodies in femoral chordotonal organs (Fig.S4). Similarly, the female sterile phenotype of *gorab^1^* was also rescued by *gorab^V266P^* and *gorab^ΔC-GTD^* but not by the other mutant transgenes (Fig. 6d). Together this indicates that the centriole duplication defect of *gorab^1^*is corrected by *gorab^V266P^*and *gorab^ΔC-GTD^*. Thus, these two mutant *gorab* alleles separate the functions of Gorab showing that ability to localize to the centrosome and fulfill centriole duplication is distinct from any function of Gorab at the Golgi. By contrast, the ΔGTD, ΔN-GTD and ΔSID Gorab mutants were not able to localize either to the Golgi or to centrosomes and could not rescue the mutant phenotypes suggesting that they have completely lost both functions.

### C-terminally tagged Gorab exerts Golgi dependent cytokinesis failure

As golgins use their C-terminal sequences to make functional interactions with Golgi membranes, we asked whether we could disrupt the localisation of full-length Gorab by blocking its C-terminus. To this end we generated fly lines ubiquitously expressing C-terminally GFP tagged Gorab from a *gorab^WT^-GFP* transgene. Males carrying a single copy of this transgene were sterile reflecting cytokinesis defects in male meiosis (Fig. 7a, b). During *Drosophila* spermatogenesis, gonial cells undergo four rounds of mitosis to form cysts of 16 primary spermatocytes that undertake meiosis to generate 64 haploid spermatids. As they differentiate into sperm, each spermatid develops a mitochondrial derivative, the Nebenkern, visible as a phase-dense sphere of similar size to the haploid nucleus. Most spermatids of *gorab^WT^-GFP* males had four nuclei and a single large Nebenkern indicative of cytokinesis failure (Fig. 7b). Immunostaining revealed that the C-terminally tagged Gorab formed string-like aggregates that overlapped more than one trans-Golgi compartment in mitotically dividing cells near the apex of the testes (Fig. 7c) and which persisted into spermatocytes (Fig. 7d). These *gorab^WT^-GFP-* expressing spermatocytes developed abnormal central spindles with irregularly shaped annilin rings having multiple protrusions in late anaphase that failed to constrict in telophase (Fig. 7d). The appearance of these mutant spermatocytes strongly resembled those observed in spermatocytes of COPI deficient flies^38^ leading us to hypothesise that expression of Gorab-GFP leads to Golgi associated defects.

**Fig.7.**
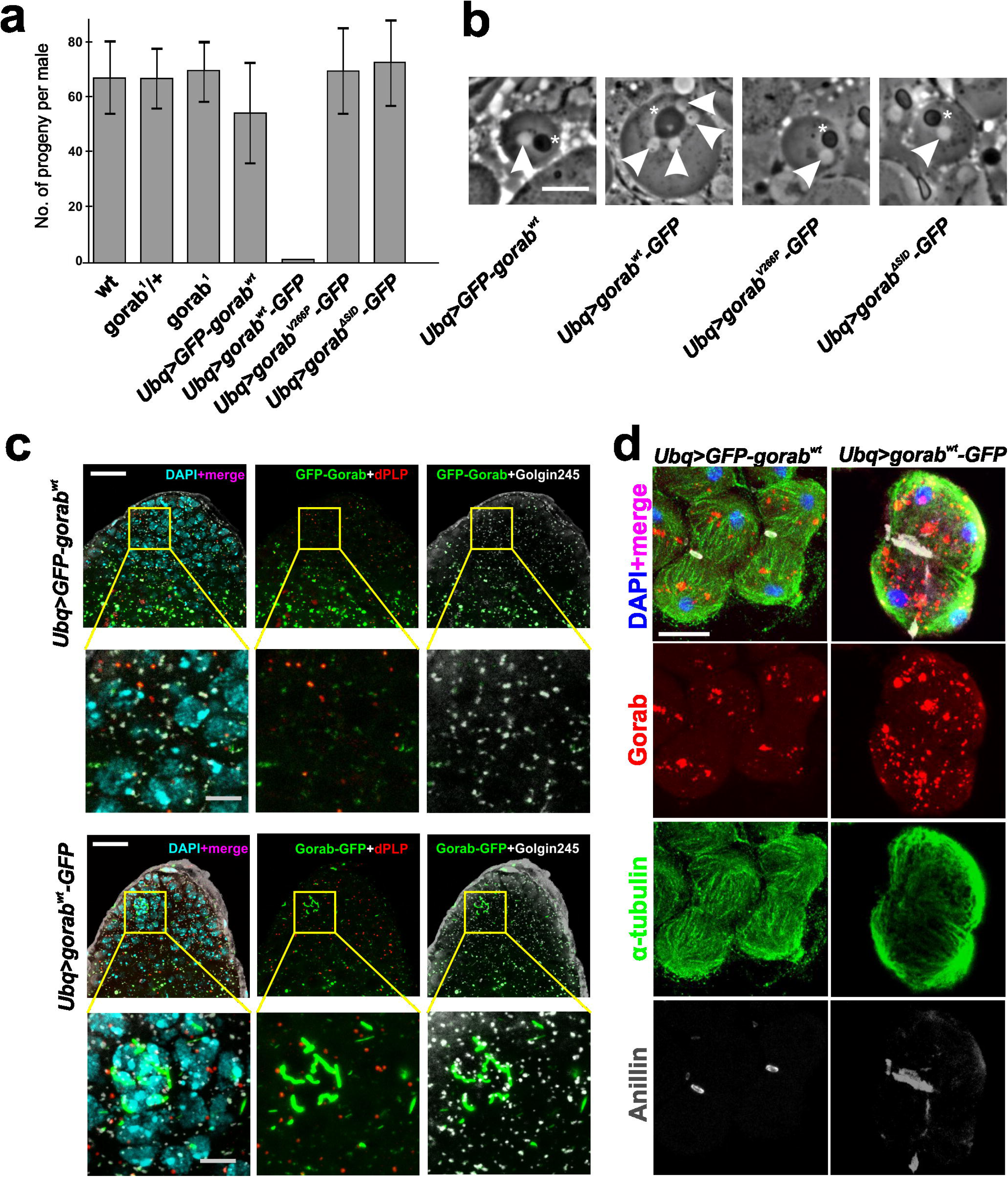
Dominant male sterility and cytokinesis defects upon expression of C-terminally GFP tagged Gorab. (**a**) Fertility of wild type, *gorab^1^* males and males expressing indicated transgenes. Individual males mated with wild type females and number of eclosed progeny recorded. Means with s.d. for 15 males. (**b**) Phase contrast micrographs of spermatids from males expressing indicated transgenes. Arrowheads indicate nuclei; asterisk, the mitochondrial derivative Nebenkern. Scale bar, 10 μm. (**c**) Apical parts of testes from males expressing N- or C-terminally GFP-tagged Gorab and stained to reveal dPLP (red) and Golgin245 (grey). Insets are 4x enlargements. Scale bars, 20 μm and 5μm (inset). (**d**) Spermatocytes in meiotic telophase from of males expressing N- or C-terminally GFP-tagged Gorab (red) and stained to reveal tubulin (green), anillin (white) and DNA (blue). Scale bar, 10 μm.

We argued that if the C-terminal tag was interfering with Gorab’s Golgi function, then inclusion of either the V266P or ΔSID mutations into a similar construct should relieve the Golgi defects that result in male sterility. Indeed, we found that transgenic flies expressing C-terminally GFP tagged V266P and ΔSID mutant versions of Gorab were fully fertile and showed no signs of cytokinesis failure (Fig. 7a, b). Thus, male sterility and cytokinesis defects observed in *gorab^WT^-GFP* flies do indeed require the C-terminally tagged protein to associate with the Golgi. Interestingly, we found that although *gorab^WT^-GFP* resulted in cytokinesis defects in male germline, it was still able to rescue the coordination and female fertility defects of *gorab^1^*mutants (Fig. S5). Thus, a C-terminal tag interferes with Gorab’s Golgi function in the male germline without affecting its function at centrioles.

## DISCUSSION

Here we provide multiple lines of evidence that unequivocally identify a tissue specific role for the Golgi-associated protein GORAB in centriole duplication. *Drosophila* Gorab forms a complex with the core component of the centriole cartwheel, Sas6. Accordingly, Gorab colocalises with Sas6 at the centriole from the very onset of procentriole formation. Centrosomes also fail to duplicate in embryos derived from *gorab* null mothers and in several tissues of *gorab* null animals, which lose coordination due to defects in the mechanosensory cilia of chordotonal organs. Basal bodies of these cilia have only a single, “mother” centriole, that has between 6 to 10 sets of microtubules, an abnormal symmetry that extends into the ciliary axoneme.

The loss of 9-fold symmetry in *gorab* mutant centrioles is strongly reminiscent of the phenotype of *Sas6* mutants ^36^. It suggests that Gorab is not only a physical partner of Sas6 required for centriole duplication but also that the partnership participates in establishing centriole symmetry. Indeed, others have reported that 9-fold symmetrical centrioles are still able to form either when the 9-fold symmetry of Sas6’s oligomerisation is destroyed ^23^or when its ability to self-oligomerize is prevented ^24^ This suggests that 9-fold symmetry is also determined by other aspects of centriole structure. Gorab could be considered to contribute, at least in part, to the provision of such unknown determinants of 9-fold symmetry. However, there is a remaining puzzle because *gorab* null mutant males are fully fertile and their spermatocytes have a correct complement of centrioles and electron microscopy showed that these all had normal 9-fold symmetry. This suggests either that maternal Gorab protein perdures at sufficient levels in male germ cells to permit centriole duplication or that Gorab function is substituted by some other component in spermatocyte centrioles. These possibilities, either of which could reflect the distinctive function and morphology of *Drosophila’s* spermatocyte centrioles, require further study.

In human cells, Gorab was first described as a Golgi associated protein ^25^. Its localisation to the trans-Golgi ^29^ is mirrored by our findings in multiple *Drosophila* tissues including salivary glands, imaginal discs, the central nervous system, and in the male and female germ lines. The association is not seen in syncytial embryos, where the Golgi apparatus has yet to form. Accordingly, Gorab’s association with components of the COPI coatomer in cultured cells, but not syncytial embryos, points to an involvement in retrograde vesicle transport from the Golgi apparatus to the ER consistent with its strong resemblance to a golgin. Golgins are rod-like proteins, which bind Rab, Arf, or ADP-ribosylation family GTPases, tethered to Golgi membranes by their C-termini and protruding outwards to capture specific vesicles at their N-termini ^39^. An overlapping specificity in their vesicle targeting provides redundancy of function. Thus, when localized to mitochondria, both golgins GMAP-210 and GM130 capture ER-derived carriers; GMAP-210 or golgin84 are able to capture vesicles containing the cis-Golgi protein ZFPL1; and so on ^40^. Such redundancy might explain why the *gorab* null mutant appears not to affect Golgi function in any obvious way. However, functional relevance of Gorab at the *Drosophila* Golgi is indicated by the cytokinesis defects in male meiosis resulting from expression of C-terminally tagged Gorab that are strikingly similar to those following disruption of COPI-mediated vesicle trafficking in spermatocytes ^38^. This accords with the association of Gorab with COPI vesicle proteins and reinforces suggestions that the functional integrity of ER and other membraneous structures is interdependent with the astral and spindle microtubules in the unusually large spindles of these cells.

However, by generating the counterpart of a missense mutant previously identified in gerodermia patients ^29^, we show Gorab’s Golgi and centriole functions to be separable. The V266P mutation prevents *Drosophila* Gorab from associating with the Golgi but is fully able to rescue the centriole duplication phenotypes of *gorab* null mutants and restore their ciliary function. Thus, although interactions of Gorab with Sas6 and the Golgi are mediated through its coiled-coil region, these requirements are not precisely overlapping. Introduction of a proline residue can have a strong influence on structure because of both the rigidity of its side chain and its ability to undergo cis-trans isomerisation. This appears to interfere with Golgi interaction without affecting ability to bind Sas6. Introducing this V266P mutation into C-terminally tagged Gorab prevents its localization to Golgi and so relieves its effect on cytokinesis. Thus the male sterility resulting from the C-terminal tag is mediated through Gorab’s Golgi association. Certainty about whether Gorab has other functions that might overlap between centriole and Golgi, which in *Drosophila* might be redundant with other Golgi proteins, must await a full characterization of Gorab’s interactions with other Golgi proteins using both genetic and molecular approaches.

Gorab appears only to be present in metazoans and so is not required for centriole duplication or Golgi function in organisms such as unicellular ciliated eukaryotes. Its appearance in the metazoans may well reflect increased proximity and functional interactions between centrosomes and cilia and the Golgi appraratus (see Introduction). The co-evolution of these organelles could have facilitated the emergence of proteins with dual functions; essentially allowing a component of one organelle to “moonlight” in its close neighbor. However, Gorab is not present in all metaozoans; it is absent, for example, from the *C.elegans* genome and so its function is not universally required. Moreover, our data indicates that even in the single species, *Drosophila,* there is a requirement for Gorab in centrioleduplication in some tissues but not others. Such tissue specificity may not be confined to flies and indeed, our present study may help clarify some recent observations about the function of the mammalian GORAB protein. A *GORAB* mutant mouse was recently generated that shows defects in hair follicle morphogenesis correlating with mild defects in Hedgehog (Hh) signaling ^41^. These mice have few primary cilia in dermal condensate cells responsible for Hh signaling in hair follicles but do have primary cilia on keratinocytes. This could be accounted for by a tissue specific failure of centriole duplication in the null mouse just as we see in null flies.

Although the above observations of defects in the development of cilia in the *GORAB* null mouse require analysis at the molecular level, they do point to a conserved role for GORAB. Both fly and human proteins are not only found at the trans-Golgi (this study; ^29^) but also in association with Sas6 at the centriole (Figs. 1,2; Fig. S6a,b). Moreover, this extends to a functional association of GORAB with Sas6 at the centriole in human cells as well as in insects (Fig. S6). Mammalian cells lacking centrosomes do not normally progress through the cell cycle as a result of a p53 dependent checkpoint pathway^24,42,43^. As loss of p53 function allows cell cycle progression to continue, we have attempted to assess the consequences of GORAB depletion upon centrosome number in a human osteosarcoma (U-2 OS) line expressing a dominant negative p53 (U-2 OS p53DD) providing a p53 compromised condition^44^. This led to cell death, possibly as a consequence of compromised Golgi function before depletion was complete, making it difficult to assess the effect upon centriole duplication. However, RNAi for GORAB enhanced the loss of centrioles seen following depletion of Sas6 alone indicative of a co-operative role of the two proteins (Fig. S6c). In a different assay, we found that depletion of human GORAB abolished the overduplication of centrosomes that occurs in the 20% of U-2 OS cells held in S-phase following treatment with Aphidicolin and Hydroxyurea A (Fig. S6d,e). Thus, as with its *Drosophila* counterpart, human GORAB is not only localised at the trans-Golgi, but is also required for centrosome duplication indicating functional homology of the fly and human proteins.

We note that the A220P mutation in humans affects GORAB’s localization to the Golgi and results in a very similar phenotype to patients with different null mutations identified by Hennies and collegues (2008) - one leading to premature termination of translation and one mutation affecting translation initiation - all of which resulted in failure to detect GORAB protein^25^. While the phenotypes resulting from all of these mutations are likely to result from defective Golgi functioning, this does not preclude an additional function of human GORAB in centriole duplication in some tissues that might be difficult to detect. In this light, it will be of future interest to re-examine the cells of patients with mutations in the *GORAB* gene for potential additional defects in centriole duplication and/or formation of primary cilia.

To conclude, our findings bring insight into the dual life of a protein with Golgi and centriole functions but also raise new future questions. An understanding of the precise role of Gorab at the Golgi will await a greater understanding of Gorab’s partners at the Golgi and how its role might be substituted by other Golgi proteins. Moreover, a full understanding of Gorab’s function in centriole duplication awaits future studies of its precise structural interactions with Sas6 and other centriole proteins and whether its role can be substituted by other proteins to allow differential functions of centrioles in different tissues.

## Acknowledgments

DMG is grateful for a Wellcome Investigator Award which supported this work. The study was initiated with support from Cancer Research UK. We would like to acknowledge Péter Deák for his encouragement and support of LK and Margit Pál for injection of CRISPR/Cas9 guide RNAs for Gorab mutagenesis, Sang Chang, Kadri Oras and Alla Madich for injection of Gorab transgenes. We also greatly appreciate the advice of Magdalena Richter and Agnieszka Fatalska in studies of protein-protein interactions. We thank Tim Megraw for GFP-Rootletin flies, Cayetano Gonzalez for YFP-Centrobin flies, Sean Munro for anti-Golgin antibodies, Kim Raj for U-2 OS^p53DD^ cell line and Janusz Debski for advice in mass-spectrometry.

